# ciliR: an R package for determining ciliary beat frequency using fast Fourier transformation

**DOI:** 10.1101/2023.12.20.572306

**Authors:** Oriane Grant, Isobel Larken, Samuel C. Reitemeier, Hannah M. Mitchison, William Dawes, Angus Phillips, Mario Cortina-Borja, Claire M. Smith

## Abstract

Cilia are motile hair-like structures that play a vital role in our body. Accurate assessment of ciliary beat frequency is pivotal for investigating ciliary dynamics and diagnosing ciliopathies. This study aims to develop software for accurately measuring the beat frequency of cilia captured using high-speed video microscopy.

To achieve this, we developed the `ciliR` package in R, which was validated against manual counting and three other automated methods of counting cilia beat frequency. The results showed that `ciliR` produced results that were comparable to manual counting. The accuracy of `ciliR` was defined by its ability to reduce noise, including only counting data in a biologically significant range (0-60 Hz).

Our software is a valuable tool for researchers in the field of ciliobiology as it offers a reliable method for detailed ciliary function analysis, thereby contributing to the broader understanding of mechanisms underlying ciliary-related disease.

We encourage researchers to try this package and feed-back their findings to the authors. Instructions for use and processes for providing feedback are provided in supplementary material.

**Summary:** ciliR is a novel R package designed for analysing ciliary beat frequency (CBF) via ImageJ and RStudio. The advantage of the ciliR system, lies in its integration with the R environment, increasing processing speed and access to data visualization tools and analysis pipelines available in other R packages. The open-source platform invites community feedback to refine functionality, aiming to advance ciliopathy research with an accessible, comprehensive toolkit.

## Introduction

Cilia are motile hair-like structures that protrude from the surface of cells and have a characteristic beating motion that helps to move fluids, particles, and mucus. In the respiratory systems, cilia help to move mucus and trapped particles towards the mouth for expulsion. In the brain, ependymal cilia lining the ventricular system serve various functions, including maintaining fluid movement, removal of debris^2^ and neuronal migration^3^. In the fallopian tube, cilia help transport ovulated eggs. Motile cilia also play a role in cell signalling and in the transport of sperm through the efferent ducts^4^ of the reproductive system. Abnormalities in cilia can cause a range of diseases, such as primary ciliary dyskinesia (Kartagener syndrome), which is characterized by respiratory problems and infertility due to the impaired movement of cilia.

Measuring cilia beat frequency (CBF) has been a topic of interest for over a century and has advanced from early methods using a stroboscope^5^ and photomultipliers^6^ to recent methods using digital cameras and high-speed video microscopy^7–11^. High-speed video microscopy enables direct observation of cilia beating and extraction of CBF data but manual counting is time-consuming and prone to human error.

To address these limitations, numerical methods that can quickly and accurately process an arbitrary number of video files provide an automated, quantitative, and unbiased solution. The basic principle behind these methods is frequency extraction from a time series of images. This process involves several steps, including segmentation of cilia in the images, reducing noise, and measurement of the cilia movement to determine their beating frequency (**Figure 1A**). Two different methods are largely used for extracting the frequency information from a data frame of pixel intensity against video frames. The first is through use of the optical density (OD) function which computes beat frequencies *f* by a translation on the time axis, *t*.^12^

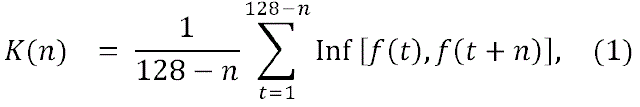

**Figure 1.**
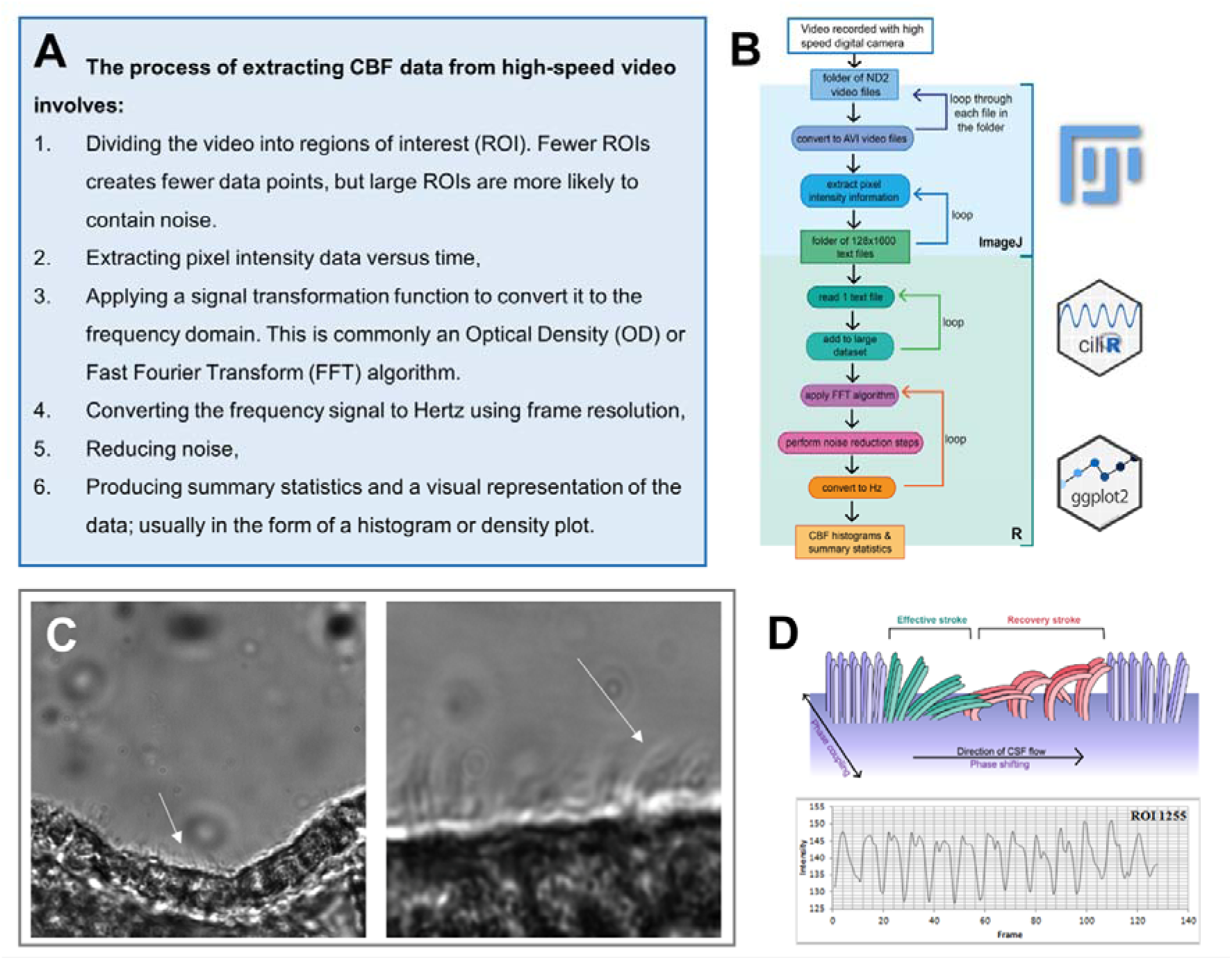
Diagrammatic view of the change in light intensity surrounding motile cilia constructed using the Volume Viewer plugin for ImageJ. (**A**) Main steps needed to calculate and analyse ciliary beat frequency from video files. (**B**) Flow chart schematic showing the processing steps using FiJi and R used to calculate ciliary beat frequency (CBF) from digital video files. (**C**) Still images from high-speed video microscopy of the mouse lateral ventricle showing 60x magnification image of the ependymal edge. White arrow is pointing to motile cilia (left panel) and cropped excerpt taken from a 60x magnification image of the ependymal edge (right pane). White arrow is pointing to motile cilia. (**D**) Ciliary beating is formed of an active effective stroke and a passive recovery stroke. Cilia in the plane of beating beat in succession; cilia in the plane perpendicular beat in unison. Below is shown how ciliary beating results in rhythmic changes in light intensity. These are extracted as pixel intensity over time/frame, which is the raw data used to obtain cilia beat frequency.

OD relies on an approximation to the average optical density for a region of interest (ROI) versus time and following^12^, we fixed as an appropriate number of frames over which the average pixel intensity for each ROI is calculated. In equation (1) is the magnitude of the translation of the OD into the time domain and is its covariance. The result of the equation is pseudo-periodic, thus only the first peak is representative of the sample’s CBF data. All other peaks are unwanted harmonics.

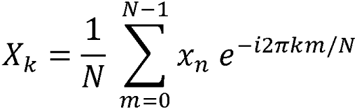

Here the impulse signal *x_n_* is translated from the time domain to a signal in the frequency domain, *X_k_*. FFT is limited to data with length equal to a power of 2 as it breaks down the function into many smaller equations of size N/2 to simplify the calculation^13^.

The available software for calculating CBF shown in Table 1 has recently been reviewed^14^. The main differences between existing tools lies in their handling of noise, which is particularly important when analysing samples with limited cilia movement. `Multi-DDM` uses the Harmonics Product Spectrum algorithm to exclude irregular frequencies as noise^15^. `ciliaFA` requires that the height of the peak in the relevant range be at least three times the size of the background noise^16^. `CiliarMove,` another open-source software for evaluating CBF has no noise reduction capability^17^. An unnamed program developed for study of primary ciliary dyskinesia (PCD) employs a time-dependent method of noise reduction by only counting ROIs with a certain number of consecutive elements– here 25 were used in the published example^19^. The output of this program includes having CBF values superimposed onto a still image of the analysed HSV, allowing the user to confirm its performance.

**Table 1.** – Methods for calculating CBF.

| Name of analysis tool | Platform | Free to use? | Reference |
| --- | --- | --- | --- |
| Manual counting | - | - | - |
| Multi-DDM | MATLAB | No* | 15 |
| ciliaFA | Image J / Excel | Yes | 16 |
| CiliarMove | Visual Studio Express (C#) | Yes | 17 |
| Sisson-Ammons Video Analysis (SAVA) | own | >3000€ | 18 |

Few studies have rigorously validated their methods as has been done before for the `ciliaFA` and `CiliarMove` programs, which reported an acceptable mean±sd difference of −0.05□±□1.25□Hz and 0.05±0.01 Hz, respectively, compared to manual counting^16,17^. The unnamed program developed by Mantovani et al. was validated using artificial models, which may have helped calibrate it but does not prove its reliability with real-world data^19^. As the `ciliaFA` program relies on a now defunct Excel 2007 language, we sought to develop and validate a new system using the R programming language, which we have named `ciliR`. A schematic for program development is shown in **Figure 1B**.

## Materials and Methods

### Image Acquisition

The methods of mounting and sectioning the mouse ependyma is described elsewhere^20^. Briefly, brain ependymal epithelium was obtained from wild type C57BL/6N mice (*n*=3). This work was performed as part of another study and <u>no additional mice</u> were used for this study. A vibratome was used to produce thin slices of the tissue whilst still able to preserve ciliary function, allowing clear images of ependymal cilia to be taken. Representative images of ciliated ependymal edges are show in **Figure 1C**. Tissue was also cultured in 35mm dishes as described^20^. The dishes were then transferred to the stage top incubation chamber (Okolab USA Inc., USA) of a Nikon Eclipse TiE microscope (Nikon Instruments Inc., Japan) with conditions set at 37°C, 5% CO_2_ and 95% humidity. The samples were explored using a 20× objective to locate the lateral ventricles. Once identified, the motility of cilia was recorded using a digital high-speed video camera (Hamamatsu C11440) with a 60× objective at a frame rate of 100-200 frames per second (fps).

### Calculation of CBF

#### Method 1: Manual Counting

As the ependymal edge is not always perpendicular in the field of view, ROIs were chosen based on where moving cilia were most easily observed by the counter. The video was played, and the counter observed 10 beat cycles of moving cilia in the ROI and noted the number of frames taken for this to occur. The following equation was then used to calculate CBF: CBF (Hz) = (frame rate/number of frames taken for 10 beat cycles to occur) × 10.

#### Method 2: ciliR

Nikon ND2 files were first converted to AVI files using a ND2 to AVI macro in FiJi (ImageJ, U.S. National Institutes of Health (ImageJ bundled with Java 18.0_172)). A second macro, named `ciliR_Pixel_Intensity_AVI.ijm,` was then used to divide each frame of an AVI file into 40×40 regions of interest (ROI) and extract the average pixel intensity for each ROI for *N*= 512 = 2^9^ frames (**Figure 2A**). The mean pixel intensities were then saved to a text file.□

**Figure 2.**
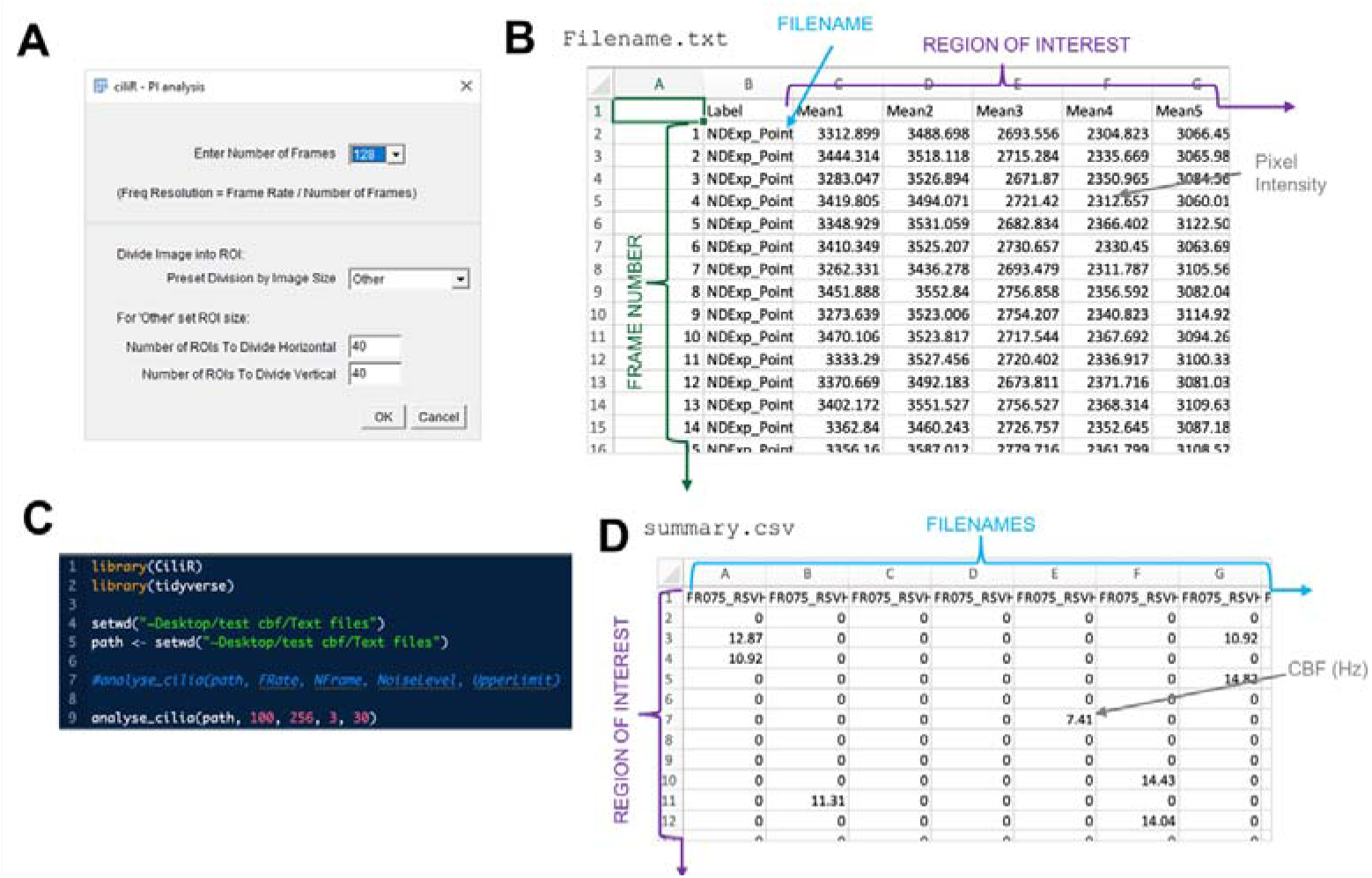
Screenshots of different files needed to complete the CBF calculation and analysis. (**A**) Screenshot of FiJi CiliR macro input message. User inputs the variables for Frame length (nearest power of 2) and number of ROIs the image is to be split into by x/y coordinates. **(B)** An example of a resulting .txt file of average pixel intensity per ROI per frame of the video. Rhythmic changes in light intensity are extracted as pixel intensity over time/frame, which is the raw data used to obtain cilia beat frequency. **(C)** Start code of `ciliR` for R showing packages needed and input variables – user must input the PATH, FRate (frame rate of recording) and NFrame (number of frames), Noise Level (default =3) and UpperLimit of CBF reading (default set at 30 Hz) **(D)** Resulting **summary.csv** file where ROI are rows, files are column and data represents the CBF in Hz.

Folders containing output files from ImageJ (**Figure 2B**) were batch processed using the package `ciliR` (ran in RStudio 1.3.959 (utilising R 4.0.2)) (see Appendix 1). The frequency signal () of each file was individually computed using an FFT algorithm. Noise reduction steps were then applied (see below) and the remaining values were converted to Hz through the following equations:

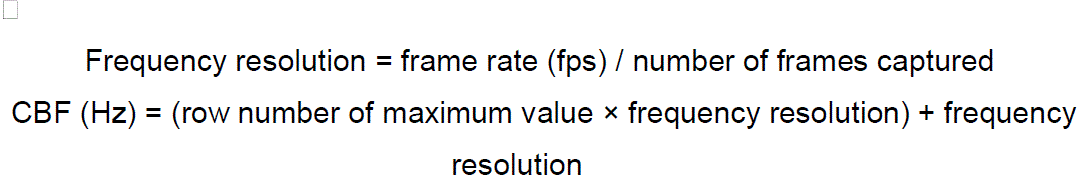

A loop was programmed so each file in the folder could be analysed in succession and stored, allowing comparative plots of each video to be generated. Summary statistics (number of ROIs, mean, mode, median, range, IQR) were collected for each file and an output graph was produced with a histogram and smoothed density estimate. Vertical lines were displayed on the graphs to indicate the position of mean and modal values.□

Several steps were taken at this stage of analysis to reduce background interference in the final output. (1) The amplitude of the peak must be 3× greater than the maximum peak of the background (defined here as a `NoiseLevel`), (2) the value for CBF must lie in a range that is clinically significant, here 3-60Hz for ependymal cilia. Data points that did not meet either criterion were removed from the plot.□A final file, combining all data from the loop provides a single data frame which facilitates visualisations using the R packages `ggplot2`^26^ and `ggridges`^27^.

#### Method 3: Multi-DDM

The software `multi-DDM` (ran in MATLAB 2020a^21^) takes ND2 video files as inputs and undergoes a three-stage analysis to produce CBF output^22^. (1) A folder of video files is converted to .MAT files through application of the multiscale differential dynamic microscopy (`multi-DDM`) algorithm: the algebraic difference between pairs of frames is calculated and a FFT analysis is performed on them to generate values in the frequency domain. The video is discretised into various numbers of boxes in the power of 2 scale (32, 64, 256, 512 and 1024) prior to applying the `multi-DDM` algorithm to them in decreasing order box size. Boxes where insufficient movement is detected (defined by the user, typically 2-3Hz) are excluded from further analysis. At this point an image of the input video can be generated showing where movement has been detected. (2) Individual MAT files are collated into one file. (3) Output histograms are produced for each individual video file. The `multi-DDM` algorithm can analyse cilia beating dynamics as well as frequency, but this was not necessary for comparison with `ciliR` ^22^.

#### Method 4: ciliaFA

The programme `ciliaFA` (ran in Image J, U.S. National Institutes of Health (ImageJ bundled with Java 1.8.0_172^23^); Microsoft Excel (2007)^24^) relies on a folder of AVI files being read and converted values of pixel intensity by ImageJ. These are then placed in Microsoft Excel, where a FFT is performed to obtain frequency values. The following noise reduction steps are used: (1) The magnitude of the frequency peak must be at least three times larger than that of the background, defined here as the maximum value of the first three FFT readings. (2) CBF values must be clinically relevant (3-60Hz for ependymal cilia). Data not fulfilling these criteria were excluded. The remaining data is inserted into an excel spreadsheet containing information on frequency peaks in each ROI, a histogram displaying all CBF values and summary statistics^16^.

### Statistical analysis

Statistical analysis was carried out using Stata/IC 16.1 for Mac (64-bit Intel) (StataCorp LLC, USA). A pairwise ANOVA was used to compare methods of calculating CBF. Bland-Altman limits of agreement were calculated from the mean difference ± 95% confidence intervals between 4803 ROIs from 3 videos processed by the `ciliR` and `ciliaFA` methods using R.

## Results

### Image acquisition

The mouse ependyma was easily identified during live microscopy. Directional fluid movement across the sample was evident and helped to locate the lumen of the ventricles under the microscope. Most cilia imaged were highly mobile and demonstrated a coordinated waveform movement.

### Comparing automatic CBF outputs

The programme `ciliR` relied on a two-step analysis. A folder of videos was first converted to text files containing information on pixel intensity using ImageJ. This folder was then processed to reduce noise, convert values into Hertz and plotted using R. A test file (file size 24.1MB, run on a 2019 MacBook with 1.3GHz Intel Core i5 processor (Apple Inc; USA) took 25.9 seconds to run through ImageJ and 2.5 seconds to be analysed in R). The main output graph for `ciliR` was a grid of histograms (see **Figure 3A)**, but via `ggplot2`^26^ and `ggridges`^27^ packages it was also capable of producing 3D density and ridge plots, as shown in **Figure 3B** and described below. 3D density plots were used to visualise where in the video the frequency signals had been detected, and in which areas the highest values resided. This allowed for confirmation that the programme had correctly identified the ependymal edge.

**Figure 3.**
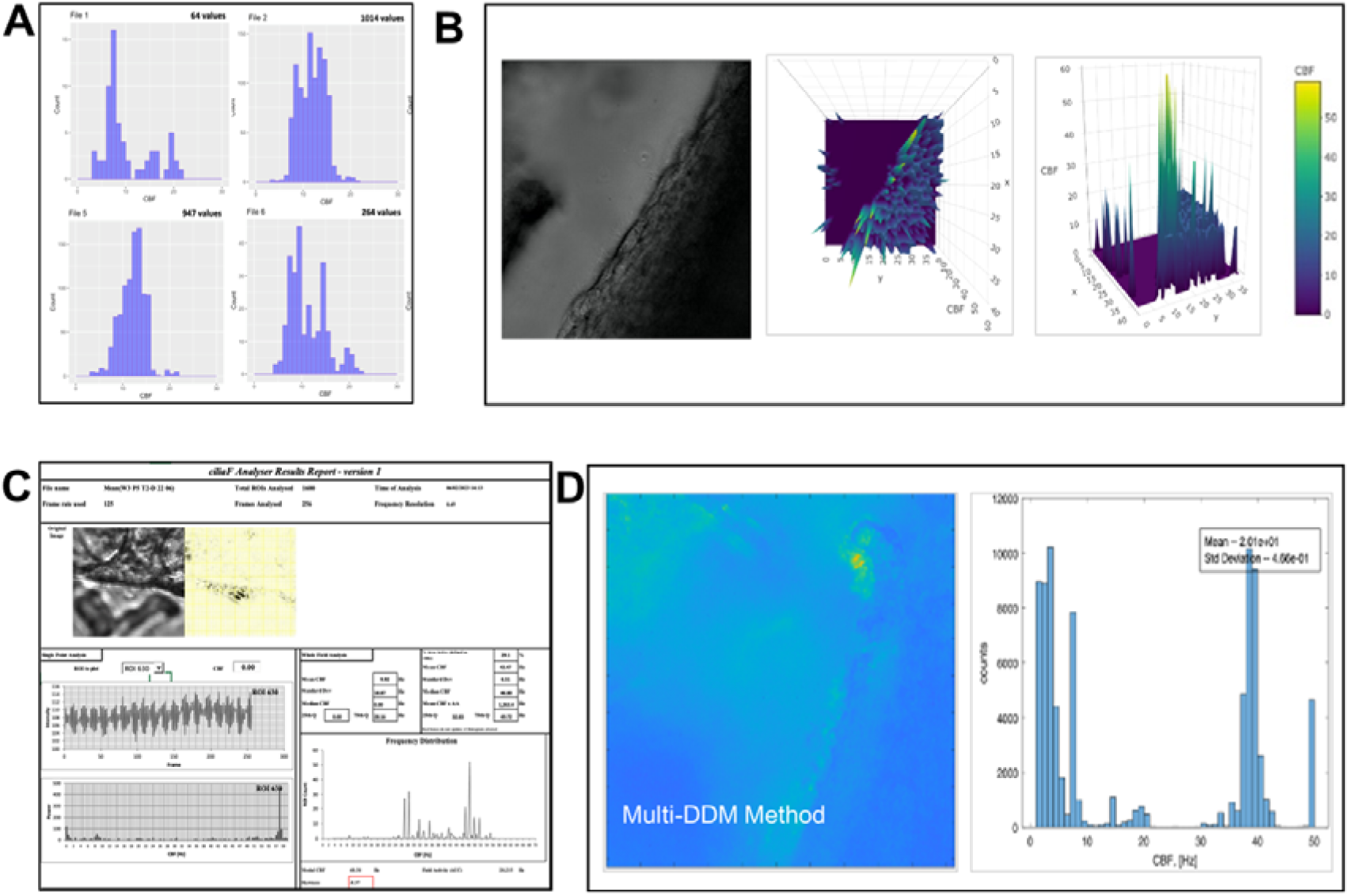
– Comparison of output graphics different automated CBF software. **(A)** Output graphics from `ciliR` (using `ggplot`). **(B)** Still image of a high-speed video of the mouse ependyma and 3D density plot showing the distribution of movement detected by `ciliR` in the video. Right panel shows a horizontal view of the same density plot. CBF ranging from 40-60Hz was detected along the ependymal edge, lower frequency signals were detected in deep tissue and an out of focus area of tissue visible to the left of the image. **(CD)** Output graphics from `ciliaFA` **(C)** and `Multi-DDM` **(D)** computer programmes.

`ciliaFA` relied on two computer programmes, but can be run from just one macro, only requiring the user to start the programme once. It produces a PDF displaying an overall histogram and summary table, as well as a histogram of CBF vs. power and a line graph of pixel intensity for a particular ROI. An example of the output PDF is shown in **Figure 3C**. The programme also yields as Excel spreadsheet containing the raw data. It took 5 minutes 40 seconds to run a single test file but was also able to batch process a series of videos.

`Multi-DDM` was by far the slowest programme of the three, taking 54 minutes to produce a data file and 2 seconds to produce output graphs. Like `ciliR`, it requires the user to start two different scripts for the different sections of the analysis. Several errors occurred when trying to graph the data through the original code, so a ‘Basic Plots’ script provided by the same developer was used instead. This created two output PDFs; a histogram showing CBF and a still image of the video that is colour coded to indicate where the package had detected cilia. These outputs are shown in **Figure 3D**. Manual counting was conducted on all videos that clearly showed cilia moving from side-to side in the image ( = 15).

T-tests were used to make pairwise comparisons between each of the three computer programs, `ciliR, ciliaFA` and `multi-DDM`, versus manual counting. Manual counting is still the most widely accepted method of assessing CBF and is therefore used as the reference point in this analysis. The normality assumption of the t-test is satisfied by the Central Limit Theorem (64 samples). There were no significant differences in the estimated CBF by manual counting (25.89 Hz; 22.81-2 8.97 Hz), `ciliaFA` (27.20 Hz; 24.28 – 30.12 Hz) and `ciliR` (27.97 Hz; 25.18 –30.77 Hz), but there was a significant difference in CBF between `multi-DDM` (11.59Hz; (9.16 – 14.03 Hz) and manual counting. A summary of the results from the methods used is shown in **Table 2**.

**Table 2.** – Comparison of methods for calculating CBF.

| Method of analysis | CBF (Hz)<br>mean (95% CI) | Time taken to process<br>1 video (s) | Software<br>used | Price (GBP)*€ |
| --- | --- | --- | --- | --- |
| Manual counting | 25.89 (22.8 - 28.9) | - | - | - |
| *ciliR* | 27.97 (25.2 - 30.8) | 25.9 (ImageJ)* | Image J | 0.00 |
|  |  | 2.5 (R)* | R | 0.00 |
| *ciliaFA* | 27.20 (24.3 - 30.1) | 340.2** | Image J | 0.00 |
|  |  |  | Excel | 119.99 |
| *Multi-DDM* | 11.59 (9.2 - 14.0) | 3240* | MATLAB | 1800.00 |

\*Run on a 2019 MacBook with 1.3GHz Intel Core i5 processor (Apple Inc; USA)
\*\*Run on a Dell OptiPlex desktop computer – *ciliaFA* requires Microsoft Excel 2007
\*Prices shown are for a personal use one-time purchase and are accurate as of 20/08/20

### Comparing CBF at a ROI level

We then performed a more detailed comparison of `ciliaFA` and `ciliR` outputs by looking at the ROI level. In total 4803 ROIs from 3 videos were processed by `ciliaFA` and `ciliR` with active cilia recorded in 2247 and 2124 ROI, respectively **Figure 4AB**. This means that 126 ROI (2.06%) were reported as active (i.e. a CBF returned) using `ciliaFA`, but the same ROIs did not past quality control using `ciliR`. These ROI returned a median CBF of 42.09 Hz (IQR ±9.56) using `ciliaFA` (**Figure 4C**). Excluding these values, the mean difference (± SD) between the methods was −2.17□±□0.25□Hz and was highly correlated (*r* =□1), meaning that if a CBF value was returned in `ciliaFA` then almost the same value was returned in `ciliR` (**Figure 4D**). There were 14 (0.66%) and 66 (3.11%) points exceeding the upper and lower limit of detection (**Figure 4E**).

**Figure 4.**
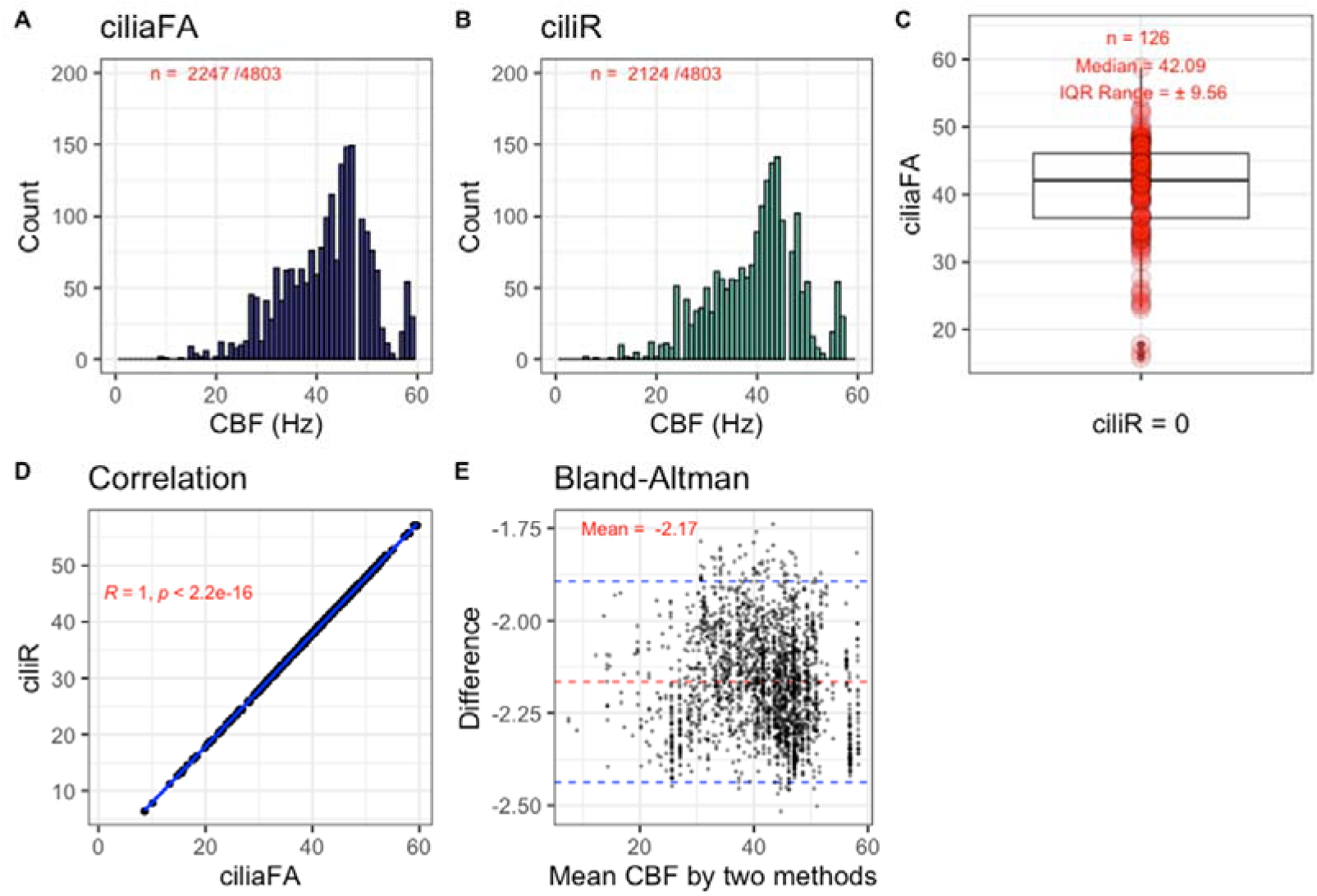
– Comparison of different automated CBF software. (ABC) Output graphics from `ciliR` (A) `Multi-DDM` (B) and `ciliaFA` (C) computer programmes. **(D)** Comparison of 2121 active ROI (total 4800) of ependymal cilia

## Discussion

We assessed the automatic readout of CBF using three methods – `ciliR, ciliaFA`, `multi-DDM` and compared these with the conventional method of manual counting. `ciliaFA` and `ciliR` both derive pixel intensity data from `ImageJ`, thus similarities were expected to be seen between these two programmes. `Multi-DDM` was selected based on the impressive graphics regarding signal decay and the degree of ciliary coordination seen in Chioccioli et al.^21^, although use of this additional analysis was not necessary for the test being carried out here.

A major advantage of `ciliR` over `ciliaFA` and `multi-DDM` is the speed of processing. Taking just 28.4 seconds compared to 340 seconds (`ciliaFA`) and 40 minutes (`multi-DDM`), `ciliR` serves a useful function in quickly returning results. All three automatic programmes can batch process a folder of video files. However, `ciliR` does require the researcher to run two programmes, first extracting information regarding pixel intensity from AVI files in ImageJ/FIJI and then importing these new text files into R to product outputs. `Multi-DDM` utilises just one application but does require the researcher to run three scripts. Furthermore, the programme was not able to be run without significant communication with the Maintainer due to numerous errors in the published script. An entirely different plotting script was eventually used due to the issues with the original code. The ability of `ciliaFA` to communicate between ImageJ and Microsoft Excel allows the researcher to leave a folder of files to be analysed overnight, which is a significant benefit.□

The 3D density plot from `ciliR` shown in **Figure 4B &C** allows for the validity of the programme to be ensured by the user, as they can confirm that values of CBF were detected in the parts of the video that contained cilia. This is of particular use when the cilia typically reside on a strip or an edge, which takes up very little of the ROIs in the image. `Multi-DDM` performs a similar function by colour coding the image based on where CBF values were extracted from as seen in **Figure 3C**, but `ciliaFA` is not capable of this step.

Both `ciliR` and `multi-DDM` are customizable programs that allow users to extensively adjust and manipulate settings to produce outputs tailored to their specific interests. This allows for a great deal of flexibility in their use. `ciliR` is the only programme of the three to run solely in freely available software, and the only programme to allow users to specify the conditions for noise reduction. MATLAB licenses are held by many universities, but personal licenses cost £1800.00 for personal use, making it poorly accessible for those without academic access. Although Microsoft Excel is a widely used application, multiple large updates to the programme over the years have made `ciliaFA` increasingly difficult to run, and previous versions such as Excel 2007, which the programme was written for, are now unable to be installed. For this study `ciliaFA` was able to be run on a computer that still possessed Excel 2007, but it did not run on more recent versions of Excel. Updates to the R programming language are typically incremental rather than overhauling, enhancing the probability that the `ciliR` program will continue to function with minimal maintenance for many years.

Although aspects such as speed, price and accessibility are important to a programme, the most fundamental feature is its ability to correctly measure cilia beating frequency. `Multi-DDM` produced significantly different results to both the method of manual counting and the two other automated methods. A reason for this likely lies in its lack of a noise reduction step to discount values *lower* than the clinically significant range, as is visible in **Figure 4E** with the high peak in CBF between 0-3 Hz. `Multi-DDM` was unable to remove this nose, leading to consistently low estimations of CBF by this programme.

Both `ciliaFA` and `ciliR` produced results that were not significantly different from manual counting. However, the results were limited by the large number of files for which `ciliaFA` could not detect any movement, or where only a very small number of ROIs made it through the rigorous noise reduction steps. While `ciliaFA` and `ciliR` generally identified similar outcomes, `ciliR` processed a significantly larger dataset and yielded results that were highly comparable to manual counting. This suggests that the improvements made to the noise reduction process in `ciliR`, as compared to `ciliaFA`, were effective.

### Implications and significance

`ciliR` serves a highly advantageous purpose by allowing high speed analysis of CBF utilising freely available software. It is a promising software for incorporating future improvements, such as further developments to include analysis of cilia beating dynamics similar to that offered by `Multi-DDM.`

### Limitations

To accommodate the variety of image capture systems, the study utilized videos recorded at three distinct frame rates: 100, 114, and 200 frames per second. The methods used to calculate CBF were able to account for this difference, but a frame rate above 200 fps significantly increases the ease of manually counting CBF and would have allowed a larger number of videos to be included in the comparison of methods aspect of this study.

## Conclusions

In conclusion, we have developed a free, fast, and reliable method for calculating CBF from HSVM imaging using R programming language. This not only provides an accurate representation of CBF in a dataset, but also extends the scope of data visualisation by utilising `ggplot2` and other R packages.

## Acknowledgments

This work was funded by grants from GOSH Children’s charity (COVID_CSmith_017), the Wellcome Trust (212516/Z/18/Z) and UKRI/ BBSRC (BB/V006738/1). This work was supported by the NIHR Great Ormond Street Hospital Biomedical Research Centre. The views expressed are those of the author(s) and not necessarily those of the NHS, the NIHR or the Department of Health. The authors would also express their gratitude to Dr Teresa Attenborough for her advice and valuable discussions that contributed to the initiation of this project.

## Author Contributions

All authors declare no conflicts of interest.

OG wrote the R code, facilitated the packaging for R, conducted experiments, analysed data, and wrote the manuscript. IL contributed to the R code. SR conducted experiments and analysed data. HM and WD provided reagents, experimental supervision, and reviewed the manuscript. AP packaged the functions. MC-B contributed to the R code, supervision of OG and conducted statistical analysis and reviewed the manuscript. CMS contributed to study conception and design, data analysis, wrote the ImageJ macro and R code and the User Instruction Guide, and contributed to the write-up of the manuscript.

