## Supplementary for "ciliR: an R package for determining ciliary beat frequency using fast Fourier transformation": CiliR package description.docx

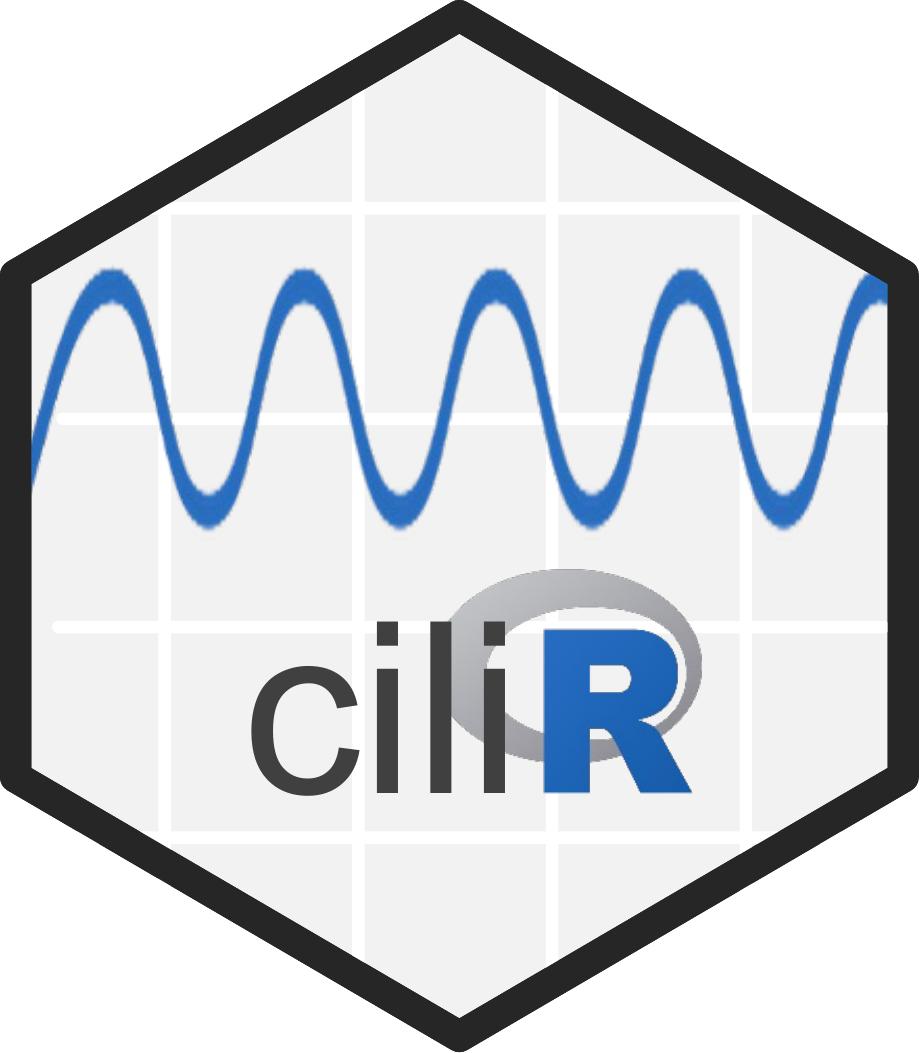
**CiliR**

```
Analyse_cilia
Description
ciliR uses a Fast Fourier Transformation to convert the pixel intensity data produced by a macro in ImageJ into cilia beating frequency (Hz)

Usage
analyse_cilia(path, FRate, NFrame, NoiseLevel, UpperLimit)


Arguments
path			Path to directory containing output from ImageJ
FRate		Frame rate used to record videos for analysis
NFrame		Number of frames being analysed
NoiseLevel	Value in Hz below which data is assumed to be noise
UpperLimit	Value in Hz describing the maximum expected value of cilia beating frequency, typically 30 for respiratory cilia and 60 for ependymal cilia

Examples
analyse_cilia(path="~/user/Documents/samples/", FRate=125, NFrame=256, NoiseLevel=3, UpperLimit=30)
```
