## Supplementary for "ciliR: an R package for determining ciliary beat frequency using fast Fourier transformation": CILIR_Pixel_Intensity_AVI.ijm

```
//To accompany the paper ciliR: a new R package using fast-Fourier transformation for determining ciliary beat frequency. 
// AUTHORS: Oriane Grant, Isobel Larken, Samuel C. Reitemeier, Hannah Mitchison, William Dawes, Mario Cortina-Borja, and Claire M. Smith. 

macro "RAW DATA FOR CILIR" {
Dialog.create("ciliaR Pixel Intensity Macro - Rules");
Dialog.addMessage("\n");
Dialog.addMessage("Before you proceed check that....");
Dialog.addMessage("1) Your image sequence file is > 128 frames in length");
Dialog.addMessage("2) Your image can be divided into at least 1600 ROI");
Dialog.addMessage("Would you like to proceed?");
Dialog.show();
// Initialize choices variables
CROP_ARRAY = newArray("128", "256", "512", "1024", "2048");
CROP2_ARRAY = newArray("0", "2", "4", "8");

//DEFAULT Properties
   XincrementInput=40;    YincrementInput=40	
// Creation of the dialog box
Dialog.create("ciliR - Pixel Intensity analysis");
Dialog.addMessage("\n");
Dialog.addChoice("Enter Number of Frames", CROP_ARRAY, "None");
Dialog.addMessage("(Freq Resolution = Frame Rate / Number of Frames)");
Dialog.addMessage("--------------------------------------------------------");
Dialog.addChoice("Enter factor required to crop to 128 frames", CROP2_ARRAY, "None");
Dialog.addMessage("--------------------------------------------------------");
Dialog.addMessage("Divide Image into ROI (Default = 1600):");
Dialog.addNumber("Number of ROIs To Divide Horizontal", XincrementInput);
Dialog.addNumber("Number of ROIs To Divide Vertical", YincrementInput);
Dialog.show();
// Feeding variables from dialog choices
CROP_TYPE=Dialog.getChoice();
CROP2_TYPE=Dialog.getChoice();
XincrementInput=Dialog.getNumber();
YincrementInput=Dialog.getNumber();
//if cropping too severe
if (CROP_TYPE=="128" && CROP2_TYPE=="4") {
			showMessage("You need at least 128F for Analysis - reduce crop increment");
			exit;
}
if (CROP_TYPE=="256" && CROP2_TYPE=="4") {
			showMessage("You need at least 128F for Analysis - reduce crop increment");
			exit;
}
if (CROP_TYPE=="128" && CROP2_TYPE=="8") {
			showMessage("You need at least 128F for Analysis - reduce crop increment");
			exit;
}
if (CROP_TYPE=="256" && CROP2_TYPE=="8") {
			showMessage("You need at least 128F for Analysis - reduce crop increment");
			exit;
}
if (CROP_TYPE=="512" && CROP2_TYPE=="8") {
			showMessage("You need at least 128F for Analysis - reduce crop increment");
			exit;
}
print("Original file size", CROP_TYPE, "Frames");
print("Crop by", CROP2_TYPE);
//set ROI 

			 Xincrement=XincrementInput;    Yincrement=YincrementInput;
	
//ETC

/////////////////////////////////////////////////////////////////////////////////////// 
//find main directory
 dir1 = getDirectory("Input Folder: contains just ND2 files to be converted"); //select an input folder.
  list = getFileList(dir1); //make a list of the filenames in the folder.
 	dir2 = dir1+" Data Files"+File.separator;
	File.makeDirectory(dir2);
		
setBatchMode(true); //turn on batch mode.
  
////////////////////////////////////////////////////////////////////////////////////////  
//Loop for cropping nd2 files
for (i=0; i<list.length; i++) {
 	showProgress(i+1, list.length);
 	filename = dir1 + list[i];
 	if (endsWith(filename, "avi")){
       open(filename);
   //    run("Bio-Formats Importer", "open=[filename] autoscale color_mode=Default crop specify_range t_begin=1 t_end=[CROP_TYPE] t_step=[CROP2_TYPE] x_coordinate_1=512 y_coordinate_1=512 width_1=1024 height_1=1024");
     //  run("Enhance Contrast...", "saturated=30 process_all");        
        
 	};   
title1 = File.nameWithoutExtension;


/////////////////////////////////////////////////////////////////////////////////////////
//PIXEL INTENSITY A
			
			run("Clear Results" , "OK");
			run("Misc...", "divide=Infinity hide");
			width=1; height=1;
			Dialog.create("Image Sampling");

			Dialog.addNumber("Width divider", Xincrement);
			Dialog.addNumber("Height divider", Yincrement);
			Dialog.addNumber("First slice", 1);
			Dialog.addNumber("Last slice", 1);
			Dialog.addNumber("Increment", 1);
			Dialog.addNumber("Substacks", 1);

			wd = Dialog.getNumber();
			hd = Dialog.getNumber();
			fs = Dialog.getNumber();
			ls = Dialog.getNumber();
			inc = Dialog.getNumber();
			substacks = Dialog.getNumber();

			getDimensions(width, height, channels, slices, frames);

					w=width/wd;
					h=height/hd;
					s=slices;
					p=fs;
					d=ls;

						for (j=0;j<hd;j++){

						for (ia=0; ia<wd; ia++) {
								x= ia*(width/wd);
								y=j*(height/hd);
								makeRectangle(x, y, w, h);

								roiManager("Add");

						};
				};

		run("Set Measurements...", "  mean display redirect=None decimal=3");
		roiManager("Show All");
		//roiManager("Save", dir1+"RoiSet.zip");
		roiManager("Multi Measure");
		selectWindow("Results");
		saveAs("Text", dir2+title1 +" Data.txt");
	
		run("Clear Results");				
	    roiManager("Select All");	
		roiManager("Delete");	
		 
run("Close All" , "OK");


	};
	

setBatchMode(false); //turn off batch mode.

showMessage("Your Files Have Been Processed And Are Ready for CiliR!");
//////////////////////////////////////////////////////////////////////////////////////////
```
