## Supplementary for "ciliR: an R package for determining ciliary beat frequency using fast Fourier transformation": User Guide for ciliR.pdf

### BETA

### User Guide for Beginners

**This guide was written by Dr Claire M. Smith, Associate Professor UCL**

Please reference all authors listed in the accompanying paper in future publications that use this software.

**TIPS ON IMAGE ACQUISITION THAT CAN IMPROVE YOUR ANALYSES:**

Limitations of any automated software depend largely on image quality and so here are some easy steps that can improve the quality of your image and your results:

1) Use the smallest pixel size available that shows sharp, clear cilia movement.

2) Use a high-power objective or crop your image so that the motile cilia fill the image. The more 'dead space' in your image the less ROI are being used for the analyses.

3) For high frequency analysis, we recommend that the microscope bulb is connected to a stable power source of at least 110-volt AC/ 60 Hz, lower voltages will encourage the bulb to flicker within the CBF range and this will enhance background noise. We also recommend that the image is not subject to down-stream processing, such as enhanced pixel gain, as this will also enhance background noise.

4) Capture at least double the number of frames to the frame rate of recording. We found that videos captured using high speed video cameras that capture at rates at least 120 frames per second with a length of 128 frames (to give a frequency resolution of 0.94) will give valid data for respiratory cilia. These settings capture an appropriate number of ciliary beat cycles to accurately average the CBF; the lower the frequency resolution (i.e. the more frames captured at this frame rate), the greater the accuracy of CBF.

### A. How to prepare video files for ciliR

A1.1 Group all your .AVI files to be analysed in one folder. Make sure this folder only contains .AVI files and no other filetype.

A1.2 Rename each .AVI file so that it has a logical name that you can easily trace and understand.

A1.3 Each file name must be unique. All .AVI files must start with a letter not a number, must be no longer than 16 characters and contain no punctuation.

### B. How to use ciliR – Part 1 – FiJi

**For conversion of image files to pixel intensity per frame.**

B1.1 Download and install FiJi from https://imagej.net/downloads

B1.2 Choose version relevant to your operating system

B1.3 Once ImageJ is installed, select "Plugins" and the "Install". Find the relevant macro from the Supplementary Folder accompanying this paper.

*If you are using .AVI files, then this will be: "**ciliR_Pixel_Intensity_AVI**".*

*If you are using another file type, select: "**ciliR_Pixel_Intensity_BioFormats**"*

B1.4 Once installed. Restart FiJi and select "**ciliR_Pixel_Intensity_AVI**" from the "**Plugins**" tab of ImageJ (the macro will appear near the bottom of the drop down menu)

Tip: Try opening your video file with FiJi first before running the macro – it's important to know that FiJi will read your file and that you can view beating cilia before proceeding with the macro.

B1.5 Once the macro is running, this window then opens to allow you to input all the information required about your files:

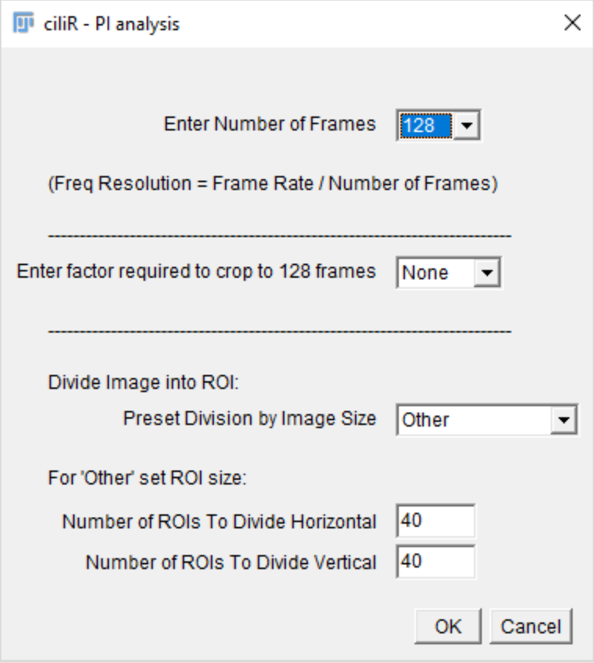


B1.6 Browse and select the folder that contains the .AVI files you want to be analysed.

B1.7 Analysis takes approximately 30s per AVI file depending on your processor speed. Do not use your computer while ciliR is running as this can interfere with processing. Wait until this message appears:

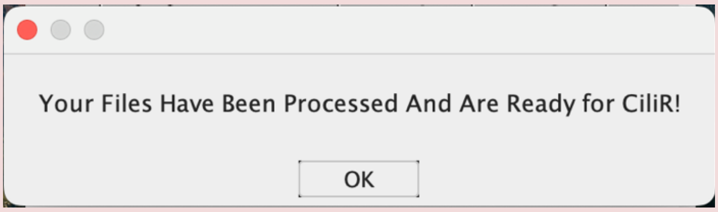


B1.8 When complete you will have a new sub-folder that contains a .txt file for each .AVI file. These files will look like this:

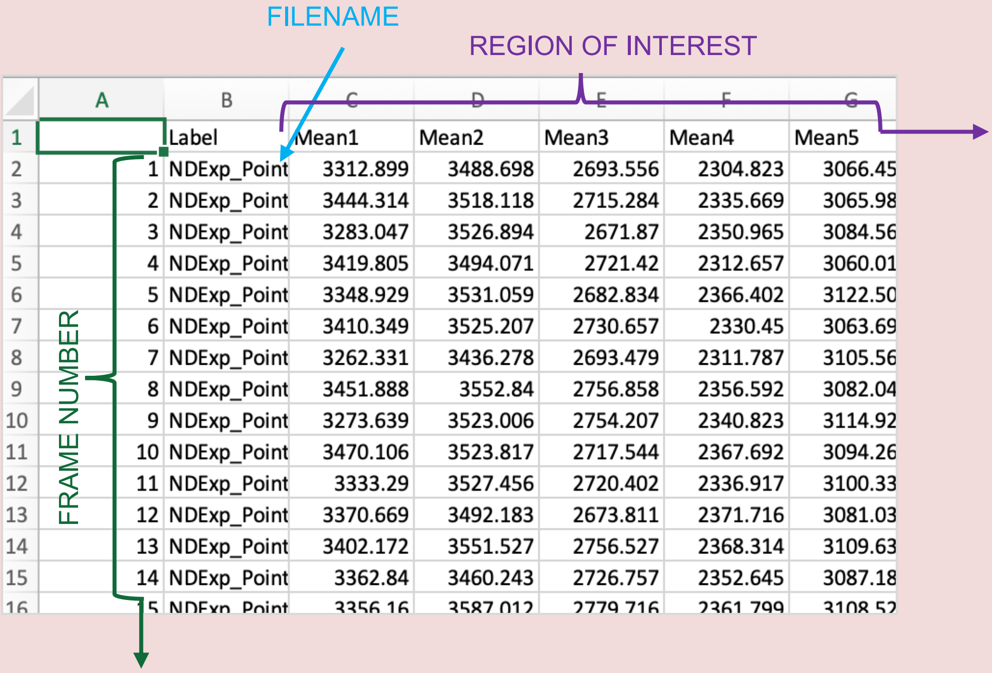


| | A | B | C | D | E | F | G |
|---|---|---|---|---|---|---|---|
| 1 | | Label | Mean1 | Mean2 | Mean3 | Mean4 | Mean5 |
| 2 | 1 | NDExp_Point | 3312.899 | 3488.698 | 2693.556 | 2304.823 | 3066.45 |
| 3 | 2 | NDExp_Point | 3444.314 | 3518.118 | 2715.284 | 2335.669 | 3065.98 |
| 4 | 3 | NDExp_Point | 3283.047 | 3526.894 | 2671.87 | 2350.965 | 3084.56 |
| 5 | 4 | NDExp_Point | 3419.805 | 3494.071 | 2721.42 | 2312.657 | 3060.01 |
| 6 | 5 | NDExp_Point | 3348.929 | 3531.059 | 2682.834 | 2366.402 | 3122.50 |
| 7 | 6 | NDExp_Point | 3410.349 | 3525.207 | 2730.657 | 2330.45 | 3063.69 |
| 8 | 7 | NDExp_Point | 3262.331 | 3436.278 | 2693.479 | 2311.787 | 3105.56 |
| 9 | 8 | NDExp_Point | 3451.888 | 3552.84 | 2756.858 | 2356.592 | 3082.04 |
| 10 | 9 | NDExp_Point | 3273.639 | 3523.006 | 2754.207 | 2340.823 | 3114.92 |
| 11 | 10 | NDExp_Point | 3470.106 | 3523.817 | 2717.544 | 2367.692 | 3094.26 |
| 12 | 11 | NDExp_Point | 3333.29 | 3527.456 | 2720.402 | 2336.917 | 3100.33 |
| 13 | 12 | NDExp_Point | 3370.669 | 3492.183 | 2673.811 | 2371.716 | 3081.03 |
| 14 | 13 | NDExp_Point | 3402.172 | 3551.527 | 2756.527 | 2368.314 | 3109.63 |
| 15 | 14 | NDExp_Point | 3362.84 | 3460.243 | 2726.757 | 2352.645 | 3087.18 |
| 16 | 15 | NDExp_Point | 3356.16 | 3587.012 | 2779.716 | 2361.799 | 3108.52 |

### C. How to use ciliR – Part 2 – R Studio

C1.1 Download and install R and R Studio from [https://rstudio-education.github.io/hopr/starting.html](https://rstudio-education.github.io/hopr/starting.html)

C1.2 Choose version relevant to your operating system

C1.3 Once R is running, install the following packages:

```
install.packages("ciliR)- via the “CiliR_1.0.tar.gz” file
install.packages("tidyverse")
```

C1.4 When you are ready to begin analysis, Open R studio and attach the “**ciliR**” package file from the “ciliR installation” folder. Then write the following lines of code in the R Script

```
library(CiliR)
library(tidyverse)

setwd("~Desktop/test cbf/Text files")
path <- setwd("~Desktop/test cbf/Text files")

#analyse_cilia(path, FRate, NFrame, NoiseLevel, UpperLimit)

analyse_cilia(path, 100, 256, 3, 30)
```

C1.5 change these parameters to those relevant to your files

setwd("path") -refers to the address of the folder containing the .txt files.

FRate - refers to the frame rate that your video files were recorded at

NFrame - refers to number of frames per video (has to = a power 2)

Code structure :
analyse_cilia(path, FRate=125, NFrame=256, NoiseLevel=3, UpperLimit=30)

Example code :
**analyse_cilia(path, 125, 256, 3, 30)**

C1.6 You can also adjust the “NoiseLevel” for noise reduction. This compares the amplitude peak of the data to the amplitude peak of the background noise. Default is 3

C1.7 Select all the code you have written.

C1.8 Click Run.

C1.9 Analysis takes approximately 3sec* per .txt file depending on your processor speed. A progress bar will appear.

When complete all your results will be saved as one dataframe referred to as "CiliaSummary" in the environment and saved as a "summary.csv" file in your working directory.

summary.csv

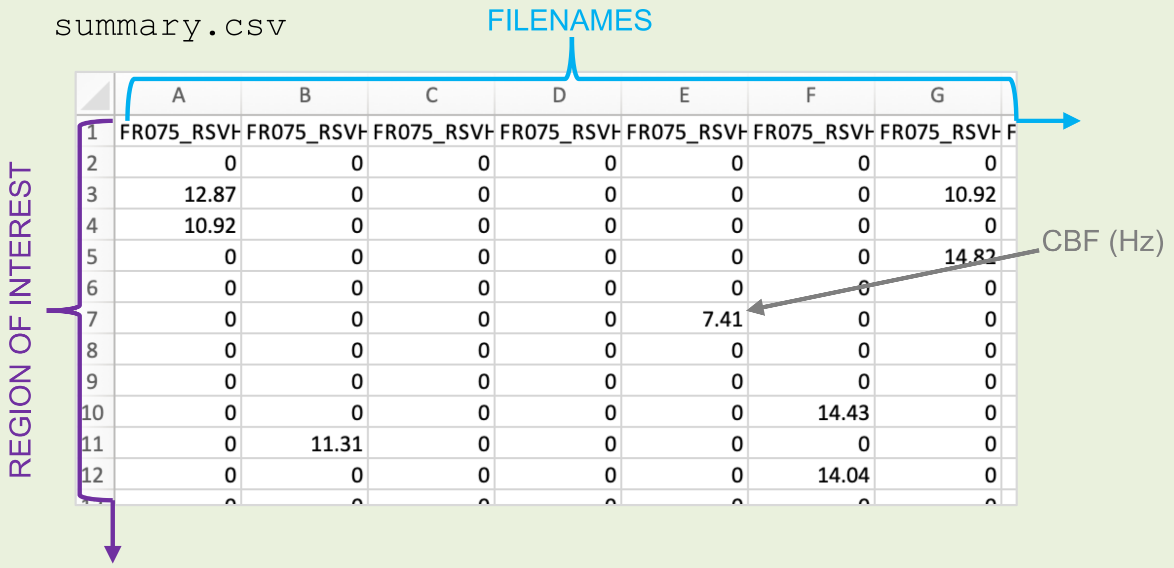


| | A | B | C | D | E | F | G |
|---|---|---|---|---|---|---|---|
| 1 | FR075_RSVH | FR075_RSVH | FR075_RSVH | FR075_RSVH | FR075_RSVH | FR075_RSVH | FR075_RSVH |
| 2 | 0 | 0 | 0 | 0 | 0 | 0 | 0 |
| 3 | 12.87 | 0 | 0 | 0 | 0 | 0 | 10.92 |
| 4 | 10.92 | 0 | 0 | 0 | 0 | 0 | 0 |
| 5 | 0 | 0 | 0 | 0 | 0 | 0 | 14.82 |
| 6 | 0 | 0 | 0 | 0 | 0 | 0 | 0 |
| 7 | 0 | 0 | 0 | 0 | 7.41 | 0 | 0 |
| 8 | 0 | 0 | 0 | 0 | 0 | 0 | 0 |
| 9 | 0 | 0 | 0 | 0 | 0 | 0 | 0 |
| 10 | 0 | 0 | 0 | 0 | 0 | 14.43 | 0 |
| 11 | 0 | 11.31 | 0 | 0 | 0 | 0 | 0 |
| 12 | 0 | 0 | 0 | 0 | 0 | 14.04 | 0 |

You can then clear objects from the workspace to free up space or continue immediately to data visualization.

* as run on a 2019 MacBook with 1.3GHz Intel Core i5 processor (Apple Inc; USA)

### 4. Data Visualization ideas

Your results will be saved as a dataframe and as a "summary.csv" file in the working directory so you can use ggplot to prepare some plots. Here is an example R code that you could use for observing the mean CBF for each file:

```
library(ggplot2)
library(reshape2)

df.long <- melt(CiliaSummary)

plot <-  df.long %>%
  ggplot(aes(x = variable, y = value)) +
  geom_boxplot()+
  labs(title = 'CBF')+
  guides(fill = guide_legend(title = "")) +
  ylab('CBF (Hz)') +
  xlab('Condition') +
  theme(legend.direction = "vertical") + theme_bw(base_size = 14)
plot
```

Or for observing the Activity of each video file:

```
df.long$Activity_group <- "Mid"
df.long$Activity_group[df.long$value >12.0] <- "Fast"
df.long$Activity_group[df.long$value <6.0] <- "Slow"

df.long <- na.omit(df.long)

grouped_data <- df.long %>%
  group_by(variable, Activity_group) %>%
  summarise(Count = n()/100, .groups = 'drop')

plot2 <-  grouped_data %>%
  ggplot(aes(x = variable, y = Count, fill = Activity_group)) +
  geom_col()+
  labs(title = 'Activity')+
  guides(fill = guide_legend(title = "")) +
  ylab('% ROI with Active Cilia')+
  xlab('Condition')+
  theme(legend.direction = "vertical") + theme_bw(base_size = 14)
plot2
```

### 5. Troubleshooting

**Common errors:**

| **Error** | **Solution** |
|---|---|
| Image J error:<br><br>Incorrect File name saved. | All .avi file names must start with a letter not a number, must be no longer than 16 characters and contain no punctuation. |
| ImageJ error:<br><br>Unable to open file | You might need to change the ImageJ macro code to make it specific to your file type. Looks at code line 78 below. Specifically, the code highlighted refers to a 3 channel RGB image where only the 3rd channel is opened to speed up processing.<br><br>`run("Bio-Formats Importer", "open=[filename] specify_range view=[Standard ImageJ] stack_order=Default c_begin=3 c_end=3 c_step=1 t_begin=1 t_end=[CROP_TYPE] t_step=[CROP2_TYPE]");` |

Any other comments or suggestions please fill in this form to get in touch with us: https://forms.office.com/e/G9sRxxTUH1

We would be happy to consider future authorship for any significant contributions to the code.
